# T-loops Refold after Telomerase Extension and Unfold Again at Late S/G2 for C-strand Fill-in

**DOI:** 10.1101/869982

**Authors:** Sabrina S.M. Mak, Phillip G. Smiraldo, Tracy T. Chow, Woodring E. Wright, Jerry W. Shay

## Abstract

Telomeres are the structures that protect the ends of each chromosome and prevent them from being recognized as broken DNA in need of repair. During human fetal development telomere length is ~15 kb and is maintained by the ribonucleoprotein telomerase. In neonates telomerase becomes spliced into inactive forms in most tissues. Afterwards, lagging strand synthesis fails to copy the very end and telomeres become progressively shorter as cells divide. Part of the mechanism for avoiding the DNA damage machinery is by forming a t-loop structure that has no free end. The telomeric 3’ overhang is inserted into the duplex DNA where it displaces the G-rich strand, forming a D-loop. It is unknown whether this D-loop is the size of the overhang, or whether branch migration results in a larger D-loop. Telomere t-loops must unfold for replication and telomerase extension during S phase. It is also unknown whether t-loops remain unfolded after replication until C-strand fill-in at late S/G2, or if they refold after replication and then unfold again for fill-in at late S/G2. We have developed a biochemical t-loop assay to address these issues, and demonstrate that most t-loops persist throughout S phase both before and after their replication.

**Highlights:**

- T-loops unfold twice during each S-phase
- Virtually all telomeres are packaged into t-loops
- Large amounts of t-loops can easily be purified without specialized materials

## INTRODUCTION

The ends of eukaryotic chromosomes are capped by repetitive sequences (TTAGGG/AATCCC) of variable length known as telomeres (1–4) with a 3’-single-stranded G-rich tail of 12-300 nucleotides (5). This 3’ overhang is believed to be inserted into the double-stranded (ds) region of the telomeres, forming a secondary lariat-like structure called a t-loop (6–9). T-loops and associated shelterin proteins protect telomeric ends from being recognized as DNA double-strand (ds) breaks, thus avoiding the activation of DNA damage repair mechanisms (10–12). T-loops have been found in *Oxytricha fallax* (9), *Trypanosoma brucei* (8), *Pisum sativum* (13), cancer cells, mouse liver cells, and normal human lymphocytes (14). Electron microscopy has demonstrated the existence of t-loops (15,16), and while t-loops are likely to be a widespread architecture for telomeres, the lack of biochemical methods to purify and analyze intact t-loops hampers progress.

In somatic cells, telomeres lose about 50-200 bps per cell division due to incomplete DNA lagging strand replication (7,17) and the inability to replicate the gap between the last priming event and the very end of the telomere (18). This phenomenon is known as the end-replication problem (19). Telomeres eventually get so short they trigger a DNA damage response and cell cycle arrest (replicative senescence) (20,21). Cancer cells avoid replicative senescence by adding TTAGGG repeats to the 3’ ends of chromosomes using the ribonucleoprotein holoenzyme, telomerase (3,22,23).

During normal human cell cycle progression, telomeric DNA is replicated throughout S phase (7,18). The overhangs after telomere lagging strand synthesis are at a nearly mature size within an hour of replication, and leading strands follow an hour later (18). T-loops would have to be unfolded for these processing events. In cancer cells telomerase extends the G-strands soon after replication, creating initial overhang sizes that are longer during S phase. The incremental C-strand fill-in that produces the final overhang takes place at the late S/G2 phase (7). We do not understand what happens to t-loops between the time a telomere is replicated/elongated in human cells and the time C-strand fill-in occurs. T-loops might re-form soon after telomere extension and unfold again at late S/G2 for C-strand fill-in, or they might stay unfolded throughout S-phase until late S/G2. This would require an unknown mechanism to prevent their recognition as damaged DNA during S-phase.

Electron microscopy (EM) imaging of t-loops requires crosslinking of ~3mg of DNA (~200-300 million HeLa cells). Crosslinking must be frequent enough to occur within the small D-loop portion of the t-loop, but the technique developed is tricky since too much crosslinking produces supercoiled balls of DNA. Only a small fraction of the t-loops are not supercoiled under conditions where some remain observable in EM. During common DNA isolation procedures, t-loops are only maintained by the hydrogen bonds between the complementary 3’ G-overhang and the strand invaded region of ds telomeres. At 55°C, branch migration of this region results in the rapid and complete unfolding of the t-loops (see figures 1 & 2A). In the present series of experiments, we have developed methods to study t-loops that only requires 5 micrograms (ug) of DNA per sample, gel electrophoresis and native in-gel hybridization of a C-rich probe to study t-loop dynamics.

**Figure 1.**
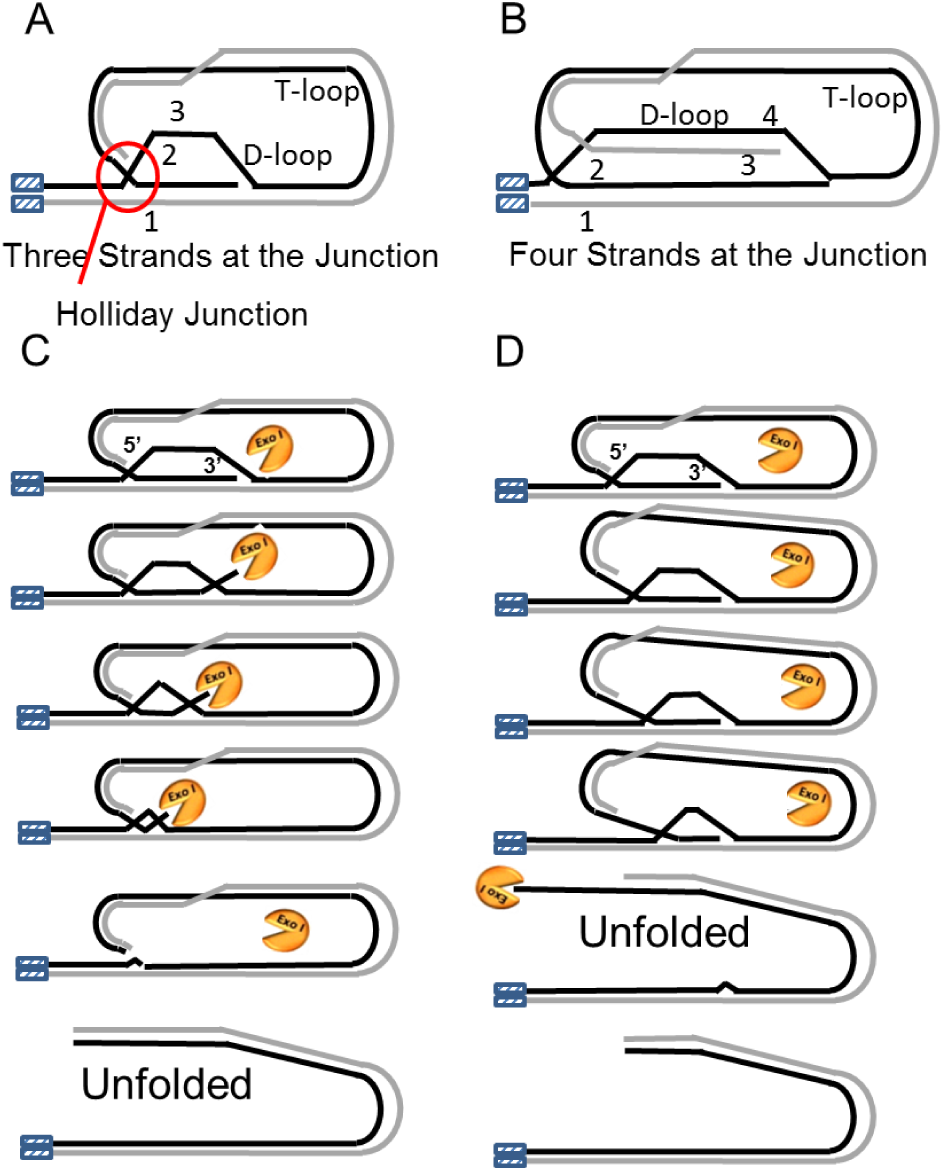
T-loop assay (t-loop vs. linear samples) A. Potential three-stranded structure fora t-loop. B. Potential four-stranded structure for a t-loop. C. In the presence of exo I, breathing of ds of the 3’ end would create a substrate for exo I digestion. This would progressively shorten the size of the overhang, thus increasing the probability of full dissociation (unfolding). D. The 3’ end is not accessible If breathing initiates at the the ds junction (the 5’ end of the overhang). The t-loop is much more stable under this condition since the overhang maintains its size.

## RESULTS

### Branch migration oligonucleotide studies provide conditions to stabilize t-loops

T-loops could exist in either 3-stranded (Fig. 1A) or 4-stranded structures (Fig. 1B). The stability of t-loops will be greater in a 4-stranded configuration, where more branch migration of both strands has to occur before dissociation. If the ds breathing happens at the 3’ end in the presence of exo I, digestion of the small ss region would produce a shorter overhang, in effect leading to a positive-feedback loop. In this case, the shorter the overhang became, the more likely it would be to lead to complete dissociation and unfolding.

Two sets of oligonucleotides with annealed ds regions and complementary single-stranded tails were combined to form a four-stranded complex (illustrated in Fig.2, sequences aligned in Fig. 3). The red and green ds regions are not homologous so that branch migration can only proceed in the direction of the black/grey sequences. The oligonucleotides contain complementary single-stranded tails of 30 nucleotides (nt), similar to the smallest HeLa and H1299 overhangs [although their average overhang is 80 nt (24)] (represented by black and gray lines in Fig. 2A and Fig. 3). Single-stranded tails of a different sequence were made to form a complex that was unable to branch migrate (Fig. 4). A lane containing the blocked complex was used to indicate the location of a four-stranded complex. The 30-nt complex was annealed at 4°C for 15 min., 60 min. and overnight (~23 hrs). The blocked complex shows four-stranded structures as early as 15 min., but it is absent in competent complexes (Fig. 2B), showing their ability to branch migrate into ds products even within 15 minutes at 4°C. Neither increasing MgCl_2_ [Fig. 2C: Mg stabilizes DNA helices (25)] or NaCl (Fig. 2D) had much of an effect at 37°C. However, including the intercalating dye ethidium bromide (EtBr) from 1-25 ug/ml stabilized the complexes even after 2 hours at 37°C and a day at 4°C (Fig 2E & F).

**Figure 2.**
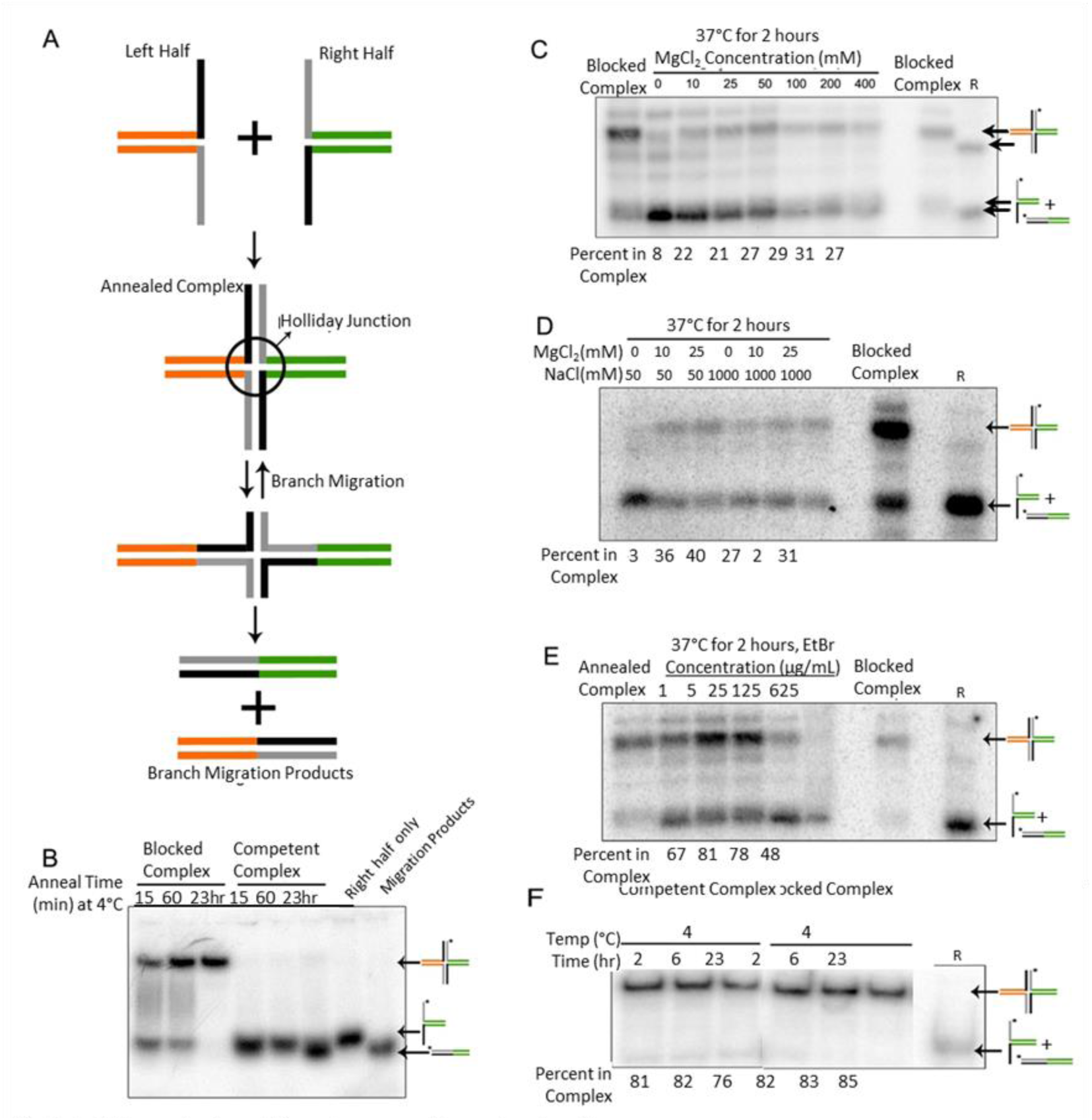
Establishment of conditions to prevent branch migration A. Schematic of the model branch migration assay. The branch migration complex were generated by first annealing oligonucleotide pairs to form left and right halves, then combining them to form a 4-stranded structure similar to a Holiday Junction. This 4-stranded structure was allowed to branch migrate (sliding of DNA back and forth) under different temperatures and salt conditions until the complete dissociation occurred that would produce two separate ds products. B. Oligonucleotides with 30 nt long tails did not stay as a complex even for 15 min at 4°C. A branch migration complex was generated with complimentary tails 30 nt long with sequences that were either branch migration competent or blocked (not complementary: see Fig. 3 for the actual sequences and how they align). The complexes were annealed at 4°C for 15 min, 60 min or 23 hrs. The blocked oligonucleotides form some 4-stranded structures within 15 minutes and they increased thereafter. None of the competent complexes stayed in 4-stranded structures and all rapidly branch migrated to the ds products. C. The effects of magnesium at 37°C were examined. Mg had little impact on complex stability. The final two lanes show size markers, where the blocked complex and the radioactively labeled right half by itself would migrate. D. The complex was annealed and allowed to branch migrate at 37°C for 2 hr in 50 or 1000mM NaCl combined with 0, 10 or 25mM MgCl_2_. Maximum stabilization occurred at 25 mM of MgCl_2_. Increasing NaCl above 50mM had no positive effect in the presence of 25 mM Mg. A blocked complex was included to visualize where the 4-stranded complex would run. R indicates where the radioactively labeled right half runs and approximately where branch migrated ds products would run. E. Ethidium bromide (EtBr) stabilizes complexes. The oligonucleotides were annealed and allowed for branch migration at 37°C for 2 hr at various concentrations of the intercalating dye ethidium bromide. 5-25 ug/mL of EtBr produced the highest amount of 4-stranded complex (~80%). 125 and 625ug/ml EtBr may have distorted the DNA enough to interfere with complex formation. F. Preservation of complexes with short tails. With salt conditions optimized plus EtBr, oligonucleotides with 30 nt tails can now be preserved as four-stranded complexes. The same oligonucleotide complexes as in B with complementary tails of 30 nts were annealed for 2, 6, and 23 hrs at 4°C under the improved conditions (50 mM of MgCl_2_, 50 mM of NaCl, and 5 ug/mL of EtBr). Almost all (~80%) of the complexes could be stabilized at 4°C even for prolonged periods of time.

**Figure 3.**
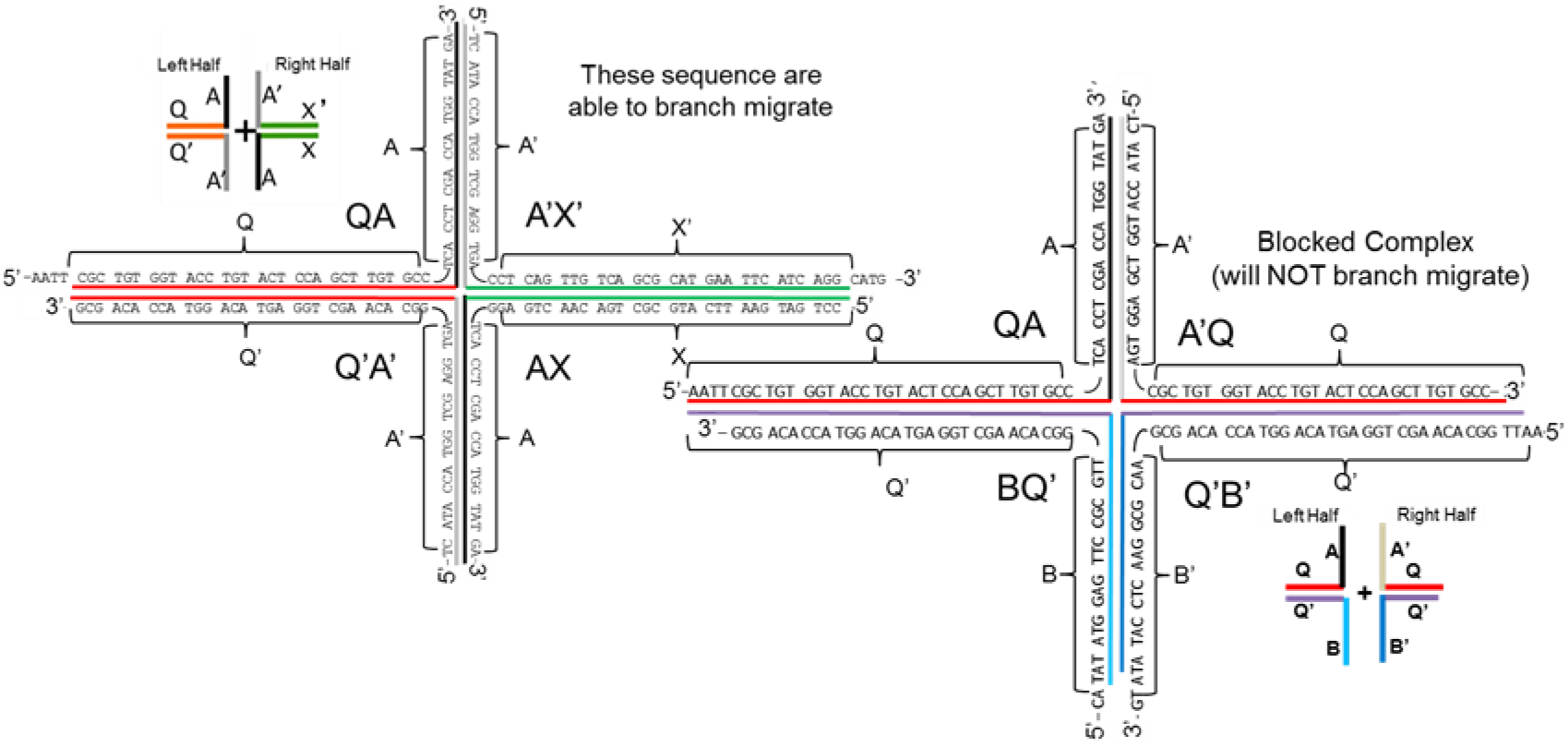
Sequence Alignment of Branch Migration Complexes. The sequences used to assemble branch migration competent and blocked complexes are shown. Pairs of sequences were first annealed in low salt to form the left and right halves, and then assembled into complete complexes at 4°C.

**Figure 4.**
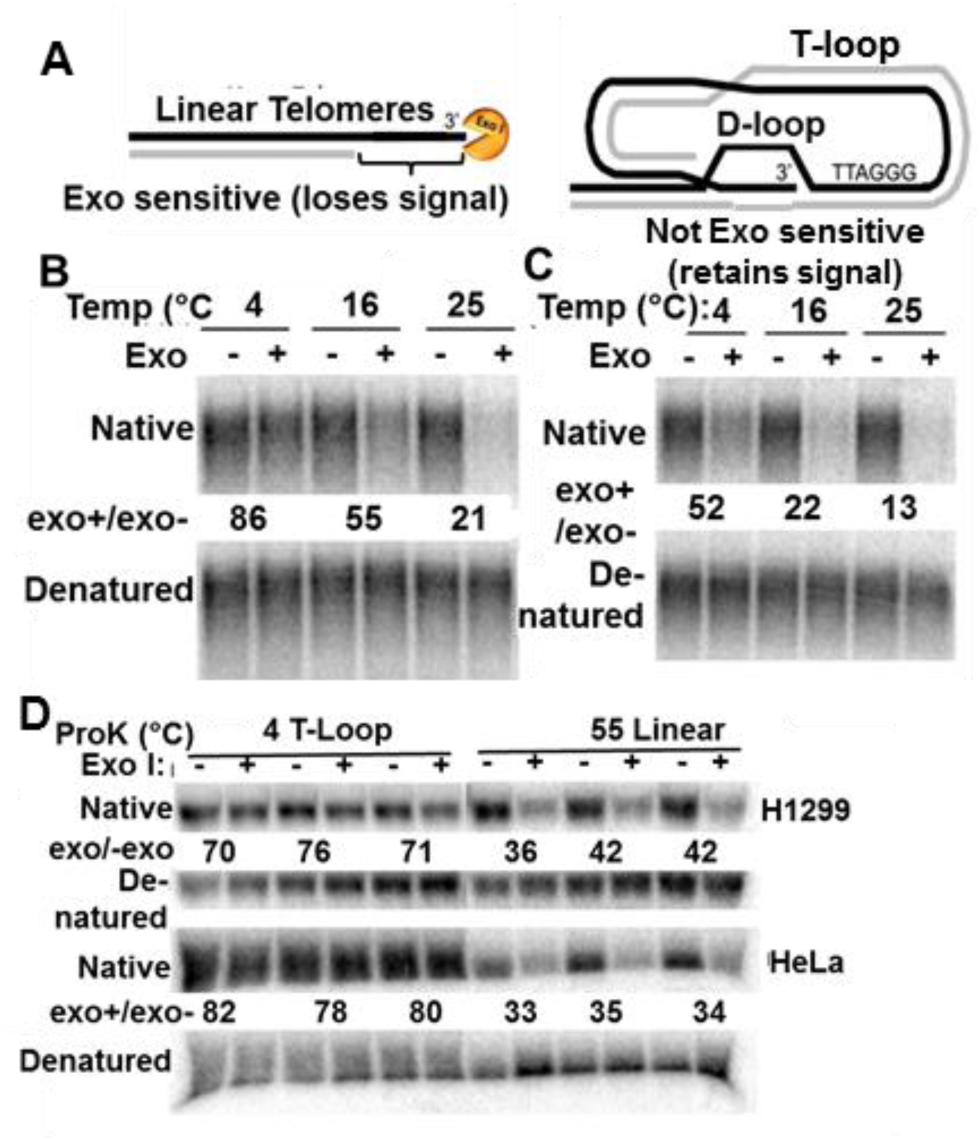
Development of the T-loop Assay. A. Schematic of Exonuclease I action at telomeres. Exonuclease I digests ss DNA in the 3’→5’ direction. In linear telomeres (left image), Exo I removes the 3’ overhang. Digestion is inhibited in t-loops (right image) since the overhang is protected after insertion in the ds region of the telomere. B. Low temperature protects t-loops. HeLa cells were harvested under t-loop conditions (but without the EtBr used in Fig. 2E), and Exo I digested at 4, 16, and 25°C Digestion at 4°C produced a small decrease in signal.. Maximal preservation would thus require the combined protective effects of both 4°C and EtBr. C. D. as in Exo I treatment of linear DNA shows a slightly decreased signal even at 4°C. 5ug DNA (HeLa cells, isolated with Quick Prep Buffer with ProK at 55°C) was digested with Exo I at 4, 16, and 25°C for 1 hr. A 50% decreased t-loop signal was seen at 4°C and becomes greater at increased temperaturesQuantitation of the experiment in B & C. E. The 3’ end of t-loops is not protected above 5U Exo I. T-loops were digested with increasing amounts of Exo I at 4°C without EtBr. Although 80% of the signal was maintained after exposure to 5U, it was significantly reduced at higher concentrations. F. T-loop preparations using high salt and EtBr protect the ss signal. Triplicate samples of DNA harvested from H1299 and HeLa cells under t-loop (ProK at 4°C with high salt) or linear conditions (ProK at 55°C) were digested with 5U of Exo I per 5 ug DNA. We saw a consistent retention of signal in (+)Exo lanes in t-loop samples (ranging from 70-80%), and a consistent decrease in signal in linear telomeric samples (ranging from 30-40%) for both cell lines.

EtBr is an intercalating dye that might stabilize DNA and prevent branch migration. Under these conditions (10 mM Mg^2+^ and 50 mM NaCl at 37°C for 2 hours) as little as 1 ug/mL of EtBr enabled 67% of the oligonucleotides to be in four-stranded complexes (Fig. 2E). The percentage plateaued at ~80% with 5-25 ug/mL of EtBr (Fig. 2E). Higher concentrations (125 and 625 ug/mL) caused a decreased signal, perhaps by disrupting complexes. We adopted 50 mM MgCl_2_, 50 mM NaCl and 5 ug/mL EtBr at 4°C for 2 hours as our standard condition for preventing branch migration (Fig. 2F).

### Development of the t-loop assay

Typical DNA isolation procedures involve deproteinization with Proteinase K at 55°C. At this temperature, the 3’ overhang should rapidly migrate and t-loops should unfold (Fig. 1B). We tested 4°C and higher salt concentrations in an attempt to stabilize ds DNA. We used LiCl instead of NaCl since Li has less of a tendency to form G-quadruplexes than Na (26). We lysed HeLa cells with 100 mM or 600 mM LiCl in “Quick prep” buffer [100 mM EDTA (to quickly inhibit DNAse activity) and 10 mM Tris pH 7.5]. Proteinase K (2mg/mL) was added with 1% Triton X-100 in place of the usual SDS (SDS precipitates at low temperatures). The lysate was incubated at either 4°C or 55°C for 1, 2, or 4 hours before inactivating Proteinase K with 2 mM PMSF. Proteinase K digested as much at 4°C as it did at 55°C in both 100 mM and 600 mM LiCl (Supplemental Figure 1). It was clear that Proteinase K was sufficiently active in the cold to remove virtually all of the proteins under these conditions.

Some DNA digestion was needed to lower the viscosity of the DNA samples to reduce sticking to the pipet tip while loading the gel. We determined the maximum salt concentration that still allowed efficient DNA digestion by Alu I at 4°C. Genomic DNA (purified from H1299 cells using the Qiagen Midi Kit) was incubated with Alu I at 4°C for 1 hr in the presence of increasing concentrations of LiCl (Supplemental Figure 1). We chose 150 mM LiCl as the highest salt concentration for efficiently digesting DNA with Alu I at 4°C.

A P^32^-labeled C-rich probe should anneal to an intact D-loop whereas there should be no ss region left after Exo1 digestion of linear telomeres (Fig. 4A). H1299 and HeLa DNA were harvested in Quick Prep buffer supplemented with Proteinase K and 600 mM LiCl at 4°C or 100mM LiCl at 55°C. After inactivating ProK with PMSF, we added 125 mM MgCl_2_ (to block the 100 mM EDTA in Quick Prep Buffer) and 150 mM Tris pH 8.9 (to neutralize the H^+^ displaced by Mg from EDTA). Exo I caused a slight reduction in the ss signal from t-loops even at 4°C (Fig. 4B), and only digested 50% of the overhang of linear telomeres (Fig. 4C). 5U exo I was sufficient to strongly diminish signals from linear telomeres (Fig. 4D) while more than 5U Exo1 per 5 ug for 1 hr caused the loss of t-loop signals (Supplemental Figure 2). Adding ethidium bromide to these conditions gave twice the signal in triplicate t-loop samples (ranging from 70-76% in H1299 and 78-82% in HeLa cells) compared to that of linear telomeres (36-42% in H1299 and 33-35% in HeLa cells: Fig. 4D). The ~35% ss signal remaining in linear DNA after exo I might represent digestion-resistant G-quadruplexes, G-knots, distortion or melting of some ds telomeric DNA after drying on a nitrocellulose or a nylon membrane. This technique nonetheless can distinguish t-looped from linear DNA using EtBr, minimal DNA (2.5 to 5 ug) and is much easier than EM.

### Validation of the t-loop assay with electron microscopy

Telomeres purified under t-loop and linear conditions were analyzed by electron microscopy (EM). Synchronized HeLa cells over-expressing hTERT were used since they have longer overhangs (~200 nts) during S-phase before C-strand fill-in (7). The longer overhangs should produce both more stable t-loops (since the amount of branch migration needed for complete dissociation would be greater) and a larger target for crosslinking. HeLa-hTERT cells were synchronized and harvested 4 hours after the release from G1/S (in the middle of S phase). The cells were divided in two, lysed with quick prep buffer, and digested with Proteinase K at either 4°C (to preserve t-loops) or 55°C (to yield linear telomeres). Both samples were then psoralen crosslinked to maintain structures throughout the subsequent steps of telomere isolation and EM imaging. Telomeres were purified using a biotinylated oligonucleotide that contained eight CCCTAA repeats followed by a linker, such that after annealing the telomere-biotinylated oligonucleotide complexes could be pulled down with streptavidin coated beads (Fig. 5A). The purified telomeres were released in low salt by incubating the samples at 60°C for 15 mins to melt off the oligonucleotide, followed by centrifuging the samples through G25 spin columns (twice) to remove any impurities that could interfere with EM. Fractions of the input, washes and final elution from the beads were Southern blotted to quantify the yield in each sample. Typical telomere purifications recovered ~10-15% after elution and ~4-8% after G25 column cleanup (Fig. 5B). Figs.5C & 5D show examples of t-loops, with white arrows pointing to the loops. Fig. 5E shows example images of telomeres from a linear preparation. A non-biased system was used to categorize the DNA samples: random numbers were assigned to each image, which were scored as t-loop, linear or not-analyzable (N/A, too tangled to score), and only afterwards identified as being from t-loop or linear telomere preparations. Telomeres were counted from two independent experiments (400 molecules each). The first experiment gave 13% t-loops and 37% linear telomeres in the t-loop sample (Fig. 5F Table 1 insert), with only 2% scored as t-loops with 76% linear telomeres in the standard 55°C preparation. The second independent experiment found 16% t-loops and 78% linear telomeres in the t-loop sample, and 1% t-loops, 95% linear telomeres in the linear preparation. This is consistent with other EM studies, provides strong evidence that the t-loop protocol successfully enriched for t-loops, and verified that using low temperatures at all processing steps in the presence of ethidium bromide enabled t-loop preservation.

**Figure 5.**
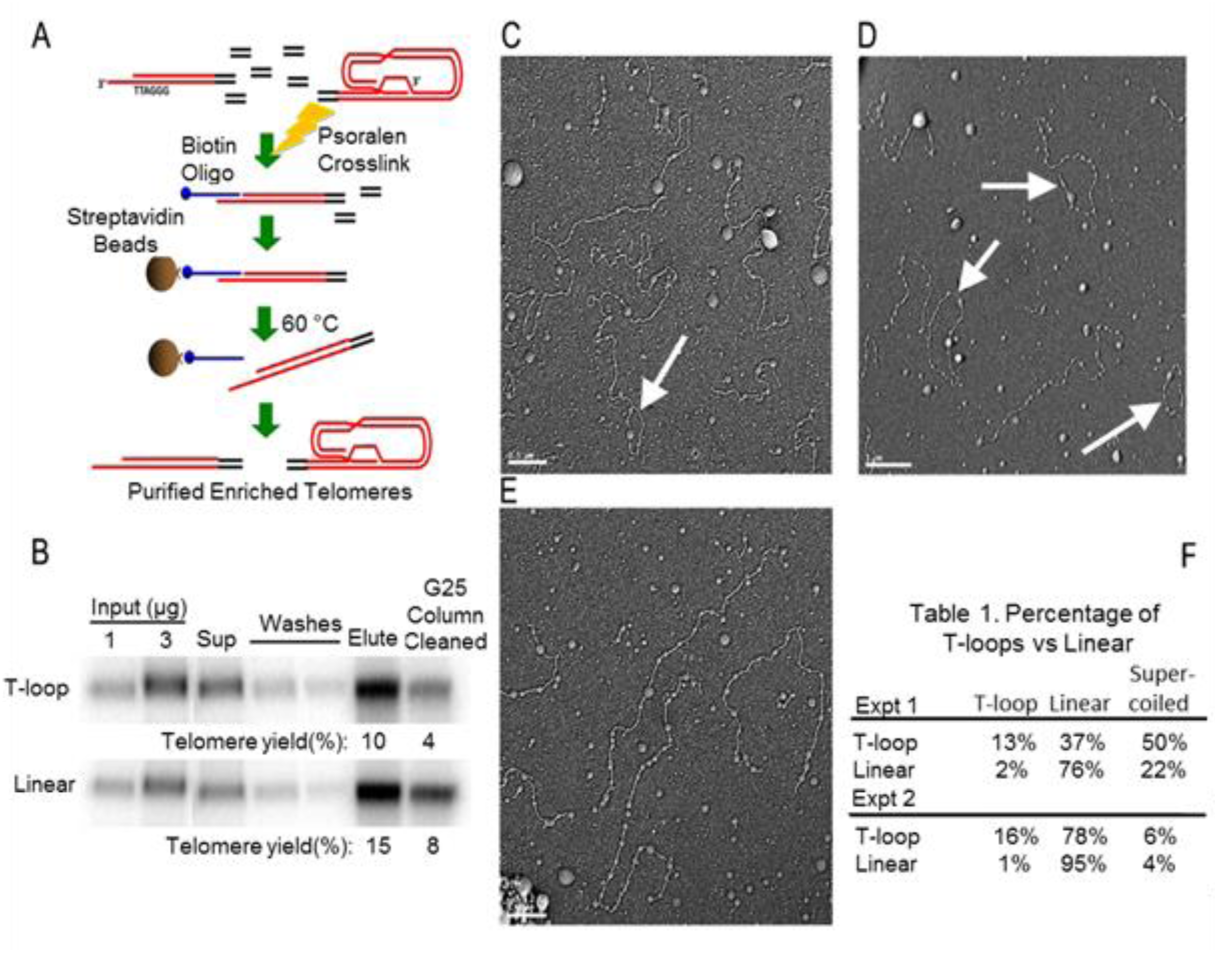
Electron microscopy validates the t-loop assay. A. Schematic of telomere purification. See Experimental Procedures for a detailed description of the use of the biotinylated oligonucleotide (that could anneal to either the ss overhang or the ss region of a D-loop) and streptavidin-coated beads to purify telomeres. B. Telomere purification at 4°C for both t-loop and linear samples gave sufficient yields for EM. HeLa-hTERT cells were harvested under t-loop conditions then psoralen crosslinked to stabilize structures during subsequent steps. Input DNA of 1 and 3ug allowed the calculation of yields. The amount of telomeres was determined from a 1:150 dilution of the supernatant after streptavidin bead capture, a 1:20 dilution of the washes, and a 1:10 dilution of both the final elution by heat and after a subsequent G25 column clean-up (to remove particulates). Typical telomere purifications yielded initial recoveries of 10-15% and 4-8% after G25 column removal of particulate contaminants. C&D. Example images of t-loops. T-loops are indicated by the white arrows. E. Example image of three linear telomeres in a linear sample. F. Quantitation of the images. Prints were numbered 1-400 for each sample type in each experiment by one individual and then scored by a second. The code was broken only after the telomeres were identified as being linear or in a t-loop. Two independent crosslinking and purification experiments were performed. There is a dramatic increase in the fraction of t-loops in the t-loop compared to the linear preparations.

### Cell cycle analysis shows t-loops refold after telomere replication

We analyzed t-loop status throughout the cell cycle. Mammalian cells replicate their telomeres throughout S-phase. T-loops must unfold for DNA polymerases to copy them. In telomerase-positive human cells, telomerase extension of the G-rich strand occurs within one hour of replication (7). T-loops must be unfolded for these processes to occur. However, C-rich strand fill-in to yield the mature overhang size is delayed until late S/G2 phase (7). If t-loops remained unfolded until the C-strands were filled-in during late S/G2 phase, some novel mechanism would have to exist for protecting them from the DNA repair machinery. Alternatively, t-loops might refold soon after replication/telomerase extension and then be unfolded again for the late S/G2 C-strand fill-in.

G1/early S synchronized HeLa-hTERT cells were released at different times in the cell cycle (0, 4, 6, 8, 10, and 12 hr after aphidocholine removal). FACs analysis showed progression through the cell cycle (Fig. 6A). DNA was isolated at 4°C in high salt as described and analyzed by Southern blotting before and after Exo I digestion. The fraction of t-loops present throughout cell cycle in the t-loop samples did not change by a significant amount (Fig. 6B). There are slightly lower ss signals (compared to the denatured DNA) in both linear and t-loop preparations at 10-12 hr as expected, since the extended overhangs should have been filled-in by then (and thus are shorter than immediately after telomerase extension. Shorter overhangs would yield less signal in linear telomeres and smaller D-loops, less stable t-loops, and lower ss signals in t-loops). We further analyzed two single time points (4-hr, Fig.6 C-E and 8-hr Fig.6 F-H) using a double-thymidine rather than an aphidocholine block (cells in S-phase when aphidocholine is added would have remained in S and would not have been synchronized at the G1/S interface, while with a thymidine block cells would finish S and arrest at the next G1/S). After 4 hr, HeLa-hTERT cells were harvested and lysed/digested according to the t-loop assay protocol. FACs confirmed cells were properly synchronized and progressing (Fig. 6C). Cesium chloride (CsCl) gradients were used to separate replicated (leading and lagging peaks were pooled) from unreplicated DNA (Fig. 6D and 6G, duplicate gradients from each experiment). The two pools were combined, precipitated and run on a gel for Southern blotting. T-loops were in the folded conformation at 4 hrs, and there is a strong overhang (D-loop) signal in the presence of Exo I (Fig. 6E). At the 8-hr time point, t-loops were again in the folded conformation since the overhang signal was preserved in the presence of Exo I (Fig. 6H, the black arrows again indicate persistence of single-stranded signal). This suggests that there is no single time point during the cell cycle (other than S/G2) where a large portion of telomeres are in the unfolded state at the same time. We conclude that t-loops in human cells are present throughout the cell cycle under normal conditions, and that individual telomeres transiently unfold and refold during S phase, so that no significant change in the amount of t-loops can be detected.

**Figure 6.**
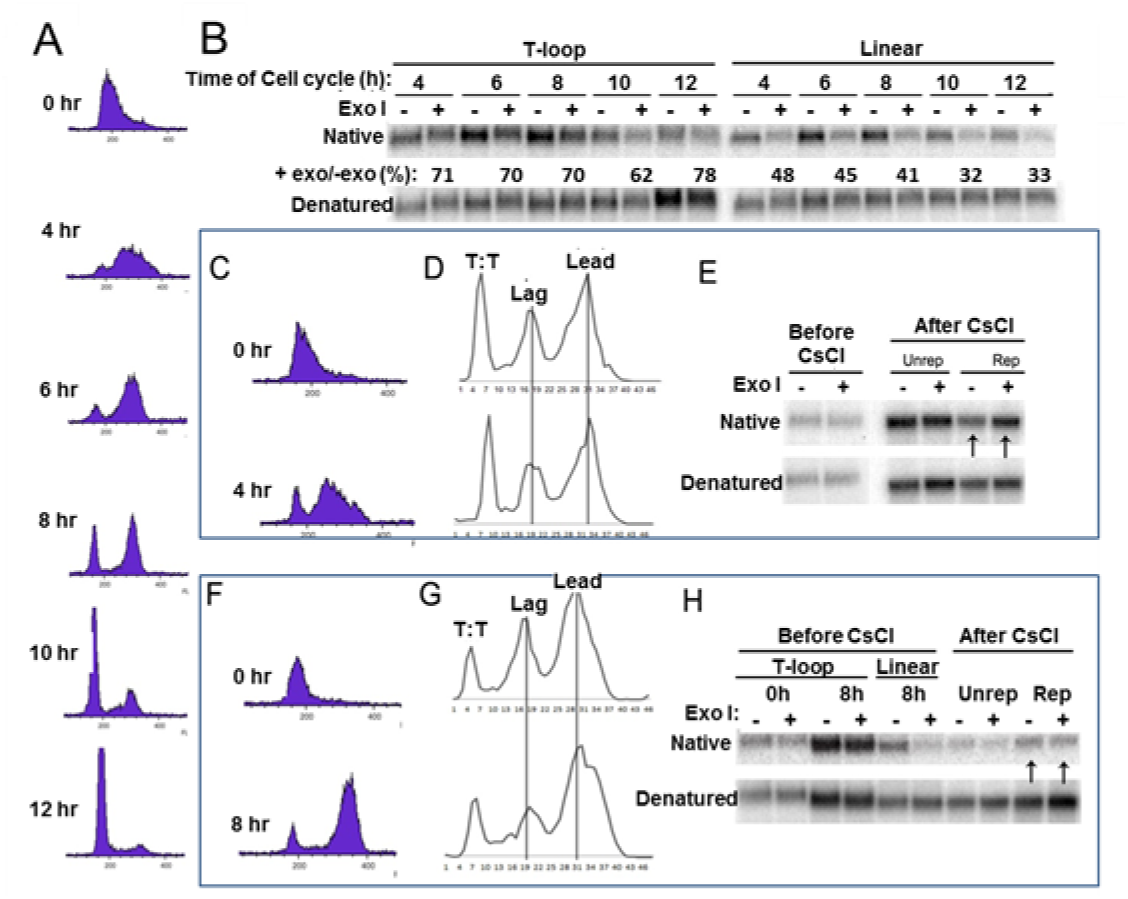
Cell cycle analysis of t-loops. A. FACS showing cells progressing throughout cell cycle. HeLa-hTERT cells, synchronized by double thymidine block, were released and followed at different time points. Aliquots of 0.5 million cells were analyzed by FACS to identify the specific phase of each cell cycle time point. B. T-loops are unfolded and refolded twice during S-phase. There was no decrease in the fraction of t-loops at any time of the cell cycle (other than at the S/G2 interface, when all of the t-loops unfold for fill-in of the extended overhangs). Each telomere replicates at a different specific time. T-loops must unfold for replication to reach the end. If telomeres remained unfolded, the exo I signal should decrease as S-phase progressed and more telomeres replicated. The maintenance of the signal suggests t-loops did not remain linear but were refolded after replication, and then unfolded again for end processing (C-strand fill-in). The denatured signal was compared to the signal from t-loop preparations to normalize for the amount of actual telomeric DNA that was loaded. One of two independent experiments is shown. C,E,E and F,G,H. FACS of HeLa-hTERT showing their cell cycle status after release for 4 or 8 hours, CsCl gradient separation of their replicated and unreplicated DNA, and sensitivity to exonuclease I. T:T represents unreplicated DNA (thymidine in both strands).The lack of sensitivity to exonuclease I (black arrows in E and H) indicates that t-loops that had replicated had been refolded and were thus resistant to digestion.

### Shorter overhangs in BJ leading strands yield less stable t-loops

In normal human BJ fibroblasts lagging strand overhangs average 100-120 nts while leading strand overhangs are only ~30-40 nts (a difference of 3-4 fold between leading and lagging overhangs) (24). We used BJ cells to determine whether the stability of t-loops was dependent on the length of the 3’ overhang (was different for leading versus lagging strands). BJ cells were labeled with IdU for 48 hours and lysed under t-loop (4°C, high salt, EtBr) or linear (55°C) conditions. Exo I was added to digest any exposed overhangs present in the samples (the t-loop assay) followed by separation of the leading and lagging strands on CsCl gradients (Fig. 7A).The lighter lagging fractions from duplicate aliquots were combined in one pool, the heavier leading strand fractions were combined in another pool, then each was precipitated and analyzed by native Southern blotting. The initial relative signal [the single-stranded signal from the t-loop preparation (before versus after Exo1 digestion) divided by the denatured signal (to correct for the amount of telomeric DNA in each sample)] was designated as 100%. Compared to this, the single-stranded signal from linearized telomere preparations was 48% (Fig. 7B, fourth lane). After CsCl gradient separation the lagging strands exhibited an increased relative signal of 73% while the leading strands showed a decrease to 39%. T-loops formed by shorter 3’ overhangs (~30-40 nts) were thus less stable in the absence of proteins than longer overhangs (~100-120 nts). This indicates that t-loops are in a 3-stranded rather than a 4-stranded configuration.

**Figure 7.**
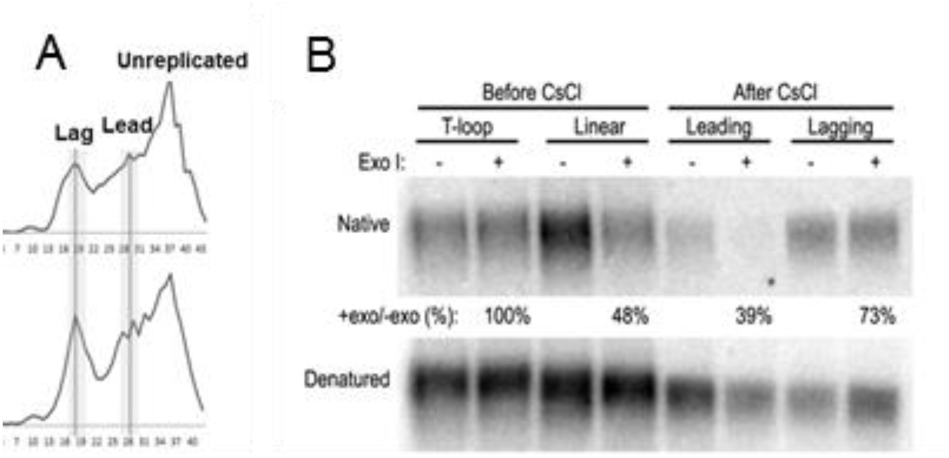
Leading overhangs (30 nt) are less stable compared with lagging overhangs (120 nt). A. CsCl gradients were used to separate leading from lagging overhangs in cells lacking telomerase activity. BJ E6/E7 cells were labeled with IdU for 48 hrs, DNA was isolated under t-loop versus linear conditions, then Exo I was added to half at 4°C for 1 hr (in the presence of EtBr) to digest overhangs. The other half was separated on CsCl gradients. Two experiments are shown. The four fractions from each peak that were pooled for analysis after CsCl are shaded in grey. B. Leading strands have less stable t-loops. Four fractions from leading and lagging peaks were pooled separately, precipitated, and tested for exo I resistance. Samples were probed with a C-rich oligonucleotide to detect the overhangs. Linear preparations (not CsCl separated) show the expected decrease in signal when treated with exo I, establishing that the exonuclease was active. After CsCl separation, leading strands from the t-loop preparations showed a dramatic decrease in the overhang signals (39% remaining) whereas lagging strands showed a preservation of the overhang signals (73% remaining), demonstrating a greater lagging strand t-loop stability. If t-loops formed a 4-stranded structure as illustrated in Fig. 1B one would expect little difference in the stability of leading vs lagging strands.

## DISCUSSION

Very little is known about t-loops since transmission electron microscopy or super-resolution light microscopy require special expertise, equipment and psoralen crosslinking. We have developed a biochemical t-loop assay that uses special salt and temperature conditions to allow intact t-loops to be isolated and analyzed.

We used oligonucleotide model systems to determine conditions to stabilize Holiday junctions. Maintaining samples at 4°C in the presence of 25 mM MgCl_2_, 50 mM NaCl, and 5 ug/mL of EtBr was effective. We later substituted LiCl for NaCl since Na^+^ promotes a greater formation of G-quadruplexes that might inhibit Exo I digestion. These conditions allowed the isolation of intact t-loops without the need for psoralen crosslinking.

We found an almost two-fold difference in the overhang signal when comparing t-loop (Pro K digestion at 4°C in high salt) vs. linear (Pro K digestion at 55°C) samples after digestion with Exo I. Increasing the enzyme concentration did not yield a higher digestion above background at 4°C, and increasing the temperature resulted in spontaneous t-loop unfolding due to increased branch migration.

In two independent EM experiments, we observed 13% and 16% t-loops using the t-loop protocol (consistent with the Griffith lab) and only 2% and 1% in linear telomere samples. This ~10-fold increase in t-loops provides strong evidence that our protocol enriched for t-loops.

Previous studies did not distinguish whether human t-loops remained unfolded after telomere replication-telomerase extension until C-strand fill-in at late S/G2 phase, or whether they refolded soon after replication and unfolded again later for the rest of the end-processing events. We demonstrate they refolded soon after replication.

BJ cells foreskin fibroblasts are known to have leading strand daughter overhangs (~30-40 nts) that are three times shorter than lagging strand overhangs (~90-120 nts) (5). Since we demonstrated that there is a large difference in the overhang signal between leading (39%) and lagging (73%) strands, these results support the interpretation that t-loops are a 3-rather than a 4-stranded structure (Fig.1A, left rather than right image).

In conclusion, our protocol establishes the dynamics of t-loops during the cell cycle and provides a method for future investigations of their function.

## METHODS

### Oligonucleotide Model Branch Migration

#### Generation of Branch Migration Complexes

DNA oligonucleotides from Integrated DNA Technologies, Inc. (IDT) are named in a 5’ to 3’ orientation, for example QA would anneal to A’Q’ in both the Q region and in the A region [forming the left half of a complex in which only the red underlined region of Fig. 3 could anneal in the initial low salt conditions due to both higher GC content (60%) and greater length (30nt vs 20nt)]. The low salt conditions prevented the shorter and 50% GC 20nt “tails” (in black or grey) from annealing]. Sequences listed below are only segments of larger oligonucleotides. For example, sequence Q is the first part of QA (Figs. 3 & 4) and the second part of A’Q (Fig. 4). Spaces are placed between appropriate nucleotides to organize them into the triplets shown in Figures 3 & 4. Pairs of oligonucleotides formed what we arbitrarily designated as a left half or a right half (Figures 2–4). Occasionally a few nucleotides were eliminated to accommodate the requirement of being able to form a 4-stranded structure without mismatches. Red, green and purple indicate the complimentary sequences within each oligonucleotide pair that are longer (30 vs 20 nt), more stable, and which can form in 10 mM Tris pH 8.0, 5 mM EDTA.

Oligonucleotide Q: 5’-AATT CGC TGT GGT ACC TGT ACT CCA GCT TGT GCC-3’

Oligonucleotide Q’: 5’-AATT GGC ACA AGC TGG AGT ACA GGT ACC ACA GCG-3’

Oligonucleotide A: 5’-TCA CCT CGA CCA TGG TAT GA-3’

Oligonucleotide A’: 5’-TC ATA CCA TGG TCG AGG TGA-3’

Oligonucleotide B: 5’-CA TAT ATG GAG TTC CGC GTT-3’

Oligonucleotide B’: 5’-AAC GCG GAA CTC CAT ATA GT-3’

Oligonucleote X: 5’-CCT GAT GAA TTC ATG CGC TGA CAA CTG AGG-3’

Oligonucleotide X’: 5’-CCT CAG TTG TCA GCG CAT GAA TTC ATC AGG CATG-3’

For each respective annealing pair, tubes containing 5 µM of each oligonucleotide in 200 µL of 10 mM Tris pH 8.0, and 5 mM EDTA were placed in a large beaker containing approximately 3.5 L of water, heated to 80°C, allowed to cool to room temperature over several hours, then stored at −20°C.

One part (volume) radioactively-labeled right half (for example, 1 µL) was mixed with three parts (volume) right half (for example, 3 µL) in 1x TNM buffer (10 mM Tris, 0.1 mM EDTA, 50 mM NaCl, 10mM MgCl_2_) and incubated at 37°C for 1 hour (see Figures 3 & 4). The relative concentration of the radioactively-labeled ligation products to the unlabeled ligation products was estimated to be approximately 1:9.

Following the 1 hour incubation, a portion of the reaction was removed, diluted approximately 20-fold in ice-cold Stop Buffer (10 mM Tris, 0.1mM EDTA, 10 mM MgCl_2_, 10% glycerol, 1ug/mL EtBr), and stored frozen at −20°C. This sample was collected for every experiment and served as a baseline for the amount of branch migration complex present at the beginning of each experiment.

#### Branch Migration Assay

A portion of the annealing reaction was diluted approximately 20-fold in 10 mM Tris, 10 mM MgCl_2_, 50 mM NaCl, 0.1 mM EDTA, pH 8 and incubated at 37°C. At specific time points (for example, 15, 30, 45, 60, 90, 120, and 180 minutes) a sample was removed, diluted approximately 3-fold in ice cold Stop Buffer (20 mM EDTA, pH 8), and stored frozen.

We examined the effects of magnesium by diluting other aliquots of the annealing reaction ~20-fold in 10 mM Tris, 10-400 mM MgCl_2_, 50 mM NaCl, and 0.1 mM EDTA. These Mg concentrations overwhelmed the chelation capacity of 0.1mM EDTA. A “no magnesium” control sample was diluted ~20-fold in 10 mM Tris, 50 mM NaCl, 10 mM MgCl_2_, and 100 mM EDTA (+100 mM EDTA = ~0 mM free MgCl_2_). These were incubated at 37°C for 2 hours, diluted approximately 3-fold in cold Stop Buffer, and stored frozen.

We studied the effects of NaCl by diluting the remaining annealed reaction ~20-fold into six aliquots containing 10 mM Tris, 50 mM versus 1 M NaCl, and 0, 10 or 25 mM MgCl_2_. These were incubated at 37°C for 2 hours, diluted approximately 3-fold in cold Stop Buffer, and stored frozen.

We determined the effects of ethidium bromide by diluting annealed samples ~20-fold in 10 mM Tris, 10 mM MgCl_2_, 50 mM NaCl, 100 mM EDTA, and 1-625 µg/mL ethidium bromide. These were incubated at 37°C for 2 hours, diluted approximately 3-fold in ice cold Stop Buffer, and stored frozen.

After 1.2% agarose electrophoresis the gels were placed on nylon membranes, dried at 60°C for approximately 2 hours, exposed to a PhosphorImager screen overnight and imaged using the Typhoon Trio Imager. The radioactive signal for the 4-way branch migration complex was normalized to the total radioactivity in the lane (ImageQuant software). The normalized value for the time=0 sample was set at 1 (or 100%) and represents the maximum 4-way complex initially present in each experiment. The normalized value for each experimental sample was compared to the time=0 sample to obtain the percent 4-way complex.

### Cell Culture

HeLa cervical carcinoma cells, H1299 lung adenocarcinoma cells, and HeLa cells overexpressing an hTERT cDNA were cultured at 37°C in 5% CO_2_ in medium containing 10% cosmic calf serum (HyClone). BJ foreskin fibroblasts expressing E6/E7 (which yielded more cells/plate than normal BJ cells following synchronization) were cultured under the same conditions but in 20% cosmic calf serum in an attempt to maximize their rate of cell division.

### T-loop Assay

HeLa cells were harvested and lysed with Quick Prep Buffer (100 mM/600 mM LiCl, 100 mM EDTA, 25% Triton X-100 and 10 mM Tris pH 7.5 (in which Tris-base had been pH adjusted with LiOH instead of NaOH to reduce possible G-quadruplex formation). 2 mg/mL of Proteinase K (Ambion) was added to ~5 ug DNA and incubated at either 4°C or 55°C for 1, 2, or 4 hrs. The reaction was stopped with the protease inhibitor phenylmethylsulfonyl fluoride (PMSF) (2 mM, Sigma-Aldrich) and loaded onto a polyacrylamide gel after heat denaturation and the addition of tracking dye. The gel was then Coomassie blue stained to estimate the protein content.

DNA from H1299 cells (using the Qiagen Blood and Cell Culture Midi Kit) was quantified using a NanoDrop DNA Spectrophotometer (Thermo Scientific). 20 U of Alu I was added to 2.5 ug of DNA in 10 mM Tris pH 7.5, 10 mM MgCl_2_, 1 mM DTT, and various concentrations of LiCl (50, 75, 150, 225, 300, 375, and 450 mM). After 1 hour at 4°C, digestion was assessed by agarose electrophoresis and UV visualization of DNA.

HeLa cells were lysed in cold Quick Prep Buffer (100 mM LiCl for linear telomeric samples and 600 mM LiCl for t-loop samples) with 25% Triton X-100. 2 mg/mL Proteinase K was added at 4°C for t-loops or 55°C for linear telomeres, and the reaction stopped with 2 mM PMSF. 125 mM of MgCl_2_ was added to neutralize the 100 mM EDTA in Quick Prep Buffer together with 150 mM Tris pH 8.9 to buffer the pH change resulting from Mg displacing the hydrogen ion bound to EDTA. 3 U Alu I per ug DNA was added to lower the viscosity of the samples. DNA was estimated as ~10 ug of DNA per 1 million HeLa cells. 0.7 U of RNase A (Qiagen) per ug of DNA was also added. After electrophoresis, a piece of agarose containing very large DNA (primarily telomeres) was then agarose dialyzed for 1 hr at 4°C (in agarose dialysis a slice of agarose is suspended in an excess of liquid containing the final desired buffer with agitation. Buffer diffuses in much more rapidly than telomeres diffuse out). For telomeres released from a biotinylated oligonucleotide, a small amount of liquid containing the telomeres was rocked over a 1% agarose plug (3-5 mm thick in a 6-well dish) to produce a buffer exchange of small molecules while the large telomeres were retained in solution. The sample was then electro-eluted in a small piece of dialysis tubing pre-coated with an excess of salmon sperm DNA, aliquoted and stored at −20°C. 20 U of Exonuclease I (NEB) was added to ~5 ug of DNA to digest free overhangs at 4°C, and the reaction stopped by adding 50 mM EDTA. These samples were then analyzed by 0.8% agarose gel electrophoresis and Southern blotting with a P32-labeled probe with the sequence (AATCCC)_4_ under native conditions. The gel was visualized with a PhoshporImager Screen on a Typhoon Trio Imager, the ss signal quantified by ImageQuant software, and the total telomere signal obtained by denaturing the gel and re-probing. This signal was used to normalize the native signal to the amount of DNA loaded.

### T-loop Cell Cycle Analysis and Electron Microscopy

Exponentially growing HeLa-hTERT cells were seeded and allowed to attach for 4 hrs before being synchronized at the beginning of S-phase with 2 mM thymidine for 18 hrs. The cells were then washed with pre-warmed PBS three times and released into pre-warmed fresh medium for 4 hrs. Aliquots of 0 hr and 4 hr after release were collected, fixed, propidium iodide (PI) stained and analyzed by FACs to confirm their cell cycle status.

DNA was isolated as in the t-loop assay and psoralen crosslinked after Proteinase K digestion. 2 U Alu I was added per ug of DNA at 4°C for 3 hr to reduce viscosity. Bead Binding Buffer (1XSSC and 0.1% Triton X-100) was added with 0.2 mg/mL of Proteinase K at 4°C for 30 mins to remove Alu I. 2 mM of PMSF was then added to inactivate Proteinase K. 5 pmol/mL of a biotinylated oligonucleotide (5’-biotin – GATCAAGCTTGAGTCGGTACC(CCCTAA)_8_-3’) called BLCTR8 [biotin-linker-C-rich telomeric repeat x8; (18)] was added at 4°C and incubated overnight to bind to the ss 3’ overhang of the telomeres (or the ss D-loop for t-loops). 16 uL of streptavidin coated beads [Dynabeads MyOne C1 (Invitrogen)], was then added to capture the biotinylated oligonuleotide-telomere complex at 4°C and incubated end-over-end overnight on a 3 RPM mechanical rotating wheel. The samples were washed by resuspending them in 200µl of 1X SSC and 0.1% Triton X-100 and then retrieving them with the Dynabeads magnet (Invitrogen), repeating using 0.2X SSC, and finally using10 mM LiCl, 1 mM Tris pH 7.5, 1 mM EDTA. Telomeres were eluted from this low salt buffer by heating the sample to 60°C for 10 mins twice in 10mM LiCl, 1mMTris pH7.5, 1mM EDTA (resuspending the beads each time). The two elutions were pooled, spun twice through illustra Microspin G-25 columns (GE Healthcare Life Sciences) to remove contaminants that might interfere with EM analysis. 1/10th of each sample was run on a 0.8% agarose gel and probed with (C_3_AATC)_4_ after being denatured to calculate the total telomere recovery using the ImageQuant software. The remainder was incubated with 250 ug/mL 4’-aminomethyl trioxalen (AMT) (Sigma-Aldrich) and crosslnked with UV at a wavelength of 365 nm.

Samples from previous step were mixed with ammonium acetate (pH 7.9) to a final concentration of 0.25 M. 4 ug/mL of cytochrome C (provided by Dr. Jack Griffith’s Laboratory) was added and allowed to sit on parafilm for 90 s. The droplet variation of the Kleinschmidt method was then used for surface-spreading DNA as described (16). Samples were visualized using an FEI Technai G2 Spirit Biotwin transmission electron microscope, and the images for publication were contrast adjusted using Adobe Photoshop. At least 400 DNA molecules were counted for each t-loop or linear telomeric sample, where random numbers were assigned to each image so that an unbiased scoring as t-loop vs. linear could be achieved.

### CsCl Gradient Separation of Leading and Lagging Daughter Telomeres

HeLa-hTERT cells were synchronized with a double-thymidine block (2 mM thymidine for 19 hr, washed with warm PBS three times, released in fresh medium for 9 hr, then incubated in 2 mM thymidine again for 17 hrs). Cells were then washed with warm PBS three times and released into warm medium containing 0.1 mM of iododeoxyuridine (IdU) for 4 or 8 hours. IdU was used in an unsuccessful attempt to get greater density separation than with BrdU. 20 U of Exo I was added per 5 ug of purified DNA and the reaction stopped with 50 mM of EDTA. Phenol, phenol:chloroform, then chloroform extractions removed Exo I and any protein contaminants still present followed by ethanol precipitation and resolubilization overnight at 37°C in 10 mM Tris (pH 7.5). CsCl gradient separation was performed as described (18). After pooling the replicated fractions and desalting via agarose dialysis followed by DNA precipitation, DNA from unreplicated and replicated fractions were run on 0.8% agarose gels and analyzed using a C-rich probe under native conditions for the t-loops present. After denaturation, reprobed signals were used to normalize the native signals. The same method was used in BJ E6/E7 cells with the exception that no synchronization was done and 0.1 mM of IdU was added for 48 hrs (instead of 24 hrs) to allow as much labeling of the replicated DNA as possible in these cells which divide much more slowly than HeLa cells and are thus difficult to synchronize.

## Acknowledgements

We thank Guido Stadler, Ph.D., and Quan Wang, Ph.D., for their assistance. We also thank Jack Griffith, Ph.D., and Ms. Smaranda Willcox for their assistance and training on visualizing DNA by EM. This research was supported by BC083222 from the Department of Defense Predoctoral Cancer Research Program (S.S.S.M.) and AG01228 from the NIH (J.W.S.). We also acknowledge the Harold C. Simmons NCI Designated Comprehensive Cancer Center Support Grant 5P30 CA142543 and the Southland Financial Corporation Distinguished Chair in Geriatric Research (J.W.S.). This work was performed in laboratories constructed with support from NIH grant C06 RR30414.

## Author Contributions

SSSM helped design the experiments, performed the bulk of the experiments and drafted the original version of the manuscript, has approved the final version and agrees to be accountable for all aspects of the paper.

PGM and TTC performed some preliminary experiments. Both have approved the final version and agree to be accountable for all aspects of the paper.

JWS helped design, interpret and edit the manuscript and agrees to be accountable for all aspects of the paper.

WEW supervised the experiments and approved the final version and agrees to be accountable for all aspects of the manuscript.

---

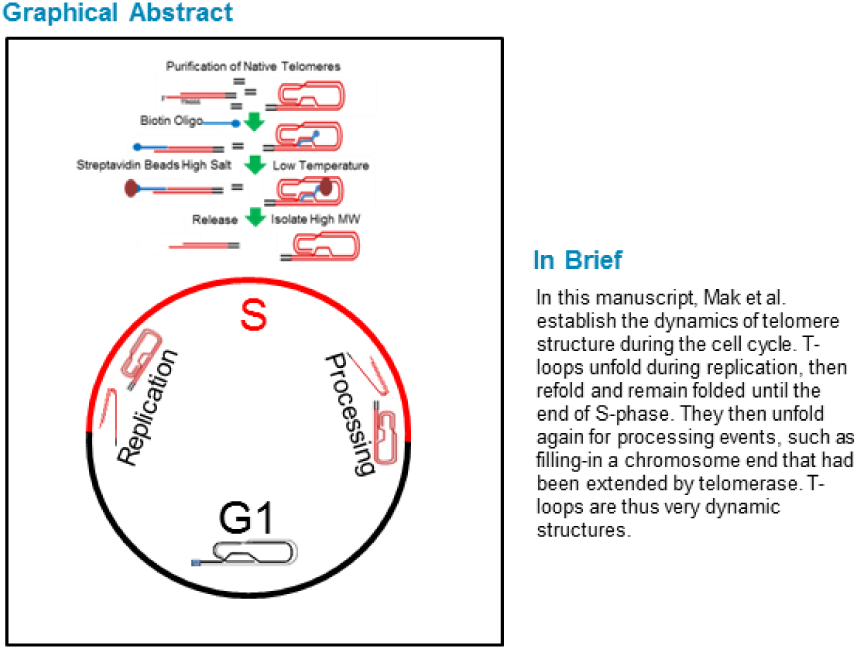

