## supplemental 2 figures for "T-loops Refold after Telomerase Extension and Unfold Again at Late S/G2 for C-strand Fill-in"

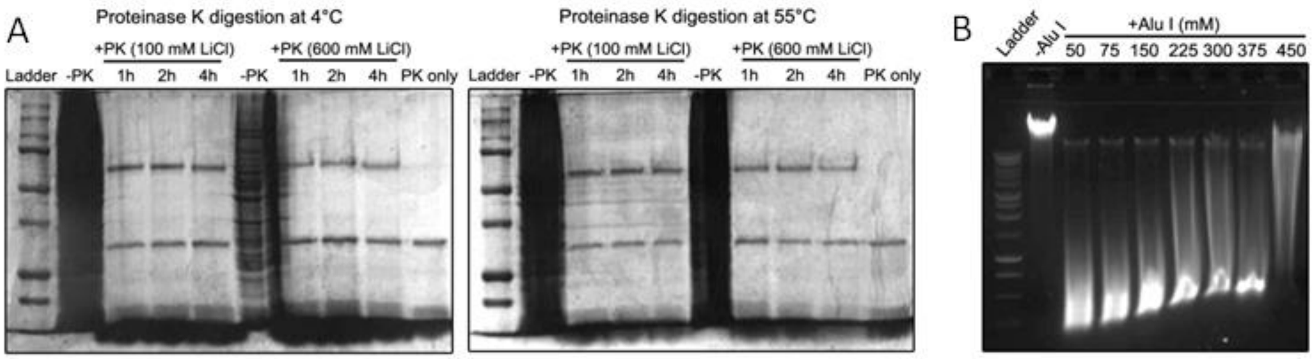


Supplemental Figure 1. Salt effects on Proteinase K and Alu I.

A. Proteinase K digests proteins efficiently at both low and high LiCl concentrations.

HeLa cells were lysed in Quick Prep Buffer with either 100 or 600 mM LiCl at 4°C and 55°C. 2 mg/mL of ProK was added for 1, 2, and 4 hours. Lanes without PK (–PK) were included to show the overwhelming amount of protein (black in silver staining) initially present in the samples. The PK only lane shows where ProK itself runs on the gel. The upper band in the +PK lanes represents an unknown PK-resistant protein. Protein digestion is efficient at both concentrations of salt at both temperatures.

B. Alu I digestion is efficient in LiCl concentrations up to 150 mM.

DNA isolated from H1299 cells using the Qiagen Blood and Cell Culture Midi Kit was digested using 20 U of Alu I per 2.5 ug of DNA at 4°C for 1 hr in the presence of increasing concentrations of LiCl. Samples were analyzed by agarose gel electrophoresis and UV visualization. Genomic DNA was well digested at up to 150 mM of LiCl, while smears of larger DNA fragments started to appear at higher concentrations.

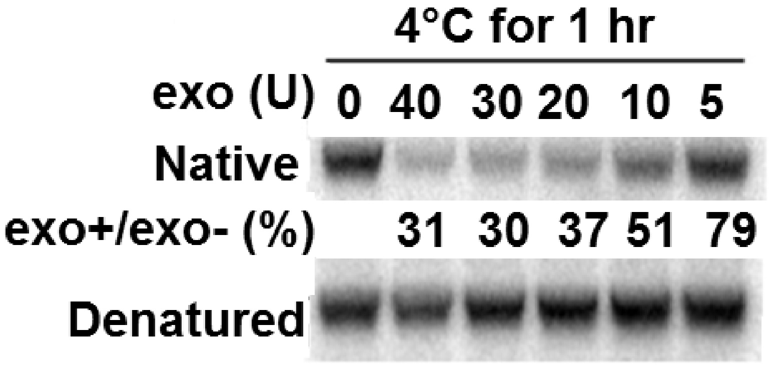


Supplemental Figure 2. . The 3’ end of t-loops is not protected above 5U Exo I. T-loops were digested with increasing amounts of Exo I at 4°C without EtBr. Although 80% of the signal was maintained after exposure to 5U, it was significantly reduced at higher concentrations
